# A tool to automate assessment of regional brain atrophy in mouse models of neurodegenerative disease

**DOI:** 10.1101/2024.11.30.626190

**Authors:** Samuel Moldenhauer, Nalini Potluri, Yuanyun Xie, Amber L Southwell

## Abstract

As life expectancy rises, so too does the prevalence of neurodegenerative diseases. Neurodegeneration causes progressive regional brain atrophy, typically initiating prior to symptom onset. Researchers measure the impact of potential treatments on atrophy in mouse models to assess their effectiveness. This is important because treatments designed to combat neuropathology are more likely to modify the disease, per contra to symptom management. Magnetic resonance imaging, while accurate in measurement of brain region structure volumes, is prohibitively expensive. Conversely, stereological volume assessment, the process of estimating the volume of individual 3D brain regions from imaged 2D brain sections, is more commonly used. This involves manually tracing brain region(s) of interest in regularly spaced imaged cross-sections to determine their 2D area, followed by application of the Cavalieri principle to estimate the volume. The pertinent caveats of this approach are the labor-intensive manual tracing process, and potential inaccuracies that arise due to human variation. To overcome these challenges, we have created a Neuropathology Assessment Tool (NAT) to automate regional brain tracing and identification using artificial intelligence (AI) and concepts from topological data analysis. The NAT was validated by comparing manual and NAT analysis of striatal volume in Huntington disease model mice. The NAT detected striatal atrophy with higher efficiency, 93.8% agreement with manual measurements, and lower inter-group variability. The NAT will increase efficiency of preclinical neuropathology assessment, allowing for a greater number of experimental therapies to be tested and facilitating drug discovery intractable neurodegenerative diseases.

## Introduction

The rising prevalence of neurodegenerative diseases such as Huntington disease (HD), coupled with the need to evaluate novel treatments, underscores the need for accurate, rapid, and resource-efficient methodologies for neuropathological assessment in animal models. While the current method of stereological volume assessment presents limitations in both efficiency and subjectivity, artificial intelligence (AI) coupled with strategies from topological data analysis (TDA) presents a promising solution.

To use stereology for regional brain volume assessment, consecutive equidistant brain cross sections spanning the region of interest are stained to reveal cytoarchitecture and imaged. A researcher then manually traces the perimeter of the region to calculate cross sectional area. The Cavalieri principle can then be applied to calculate the 3D volume using the sum of cross-sectional area and the intersection interval^1^. However, due to caveats arising from manual labor-intensive tracing within this approach, we are applying a more recent advancement in AI called self-attention, and our personal advancements upon the technique, along with our own algorithm based on TDA to not only automate but also improve the precision of stereological volumetric assessments.

Our immediate focus is to use our program to assess striatal atrophy in HD model mice. The focus on the striatum stems from observation that medium spiny neurons in the striatum, which make up the majority of its composition, are the first to undergo cell-death^2^ in HD brain. This results in progressive reduced striatal volume in both premanifest HD individuals and HD patients in comparison to control subjects. This regional brain atrophy is consistent in HD model mice^3^, this reinforces the importance of performing neuropathological assessment in the striatum when evaluating the effectiveness of potential therapies. This will allow comparisons of the method to historical data for stringent validation. Our current approach could be rapidly applied to other regions of interest, including frontal cortex and corpus callosum. These regions also exhibit progressive atrophy in HD patients in comparison to healthy control individuals^4^, which is also consistent with HD model mice, warranting our need to perform neuropathological assessments in these regions as well. Our program could lead to more rapid and accurate preclinical evaluation of potential therapies requiring fewer mice and reduced labor. Moreover, applying our technology would also lead to decreased spending on resources that would have previously been required for stereological volume assessment. It is also important to note that in addition to its utility in HD research, our program could be adapted to the study of other neurological diseases.

## Methods

### Creation of the dataset

Both Hu97/18 humanized HD model mice^3^ and Q175FDN HD model mice^5^ were aged to 12 months and perfused intracardially. Brains were post-fixed overnight in 4% paraformaldehyde, cryoprotected in 30% sucrose, and cut into 25 µM free-floating sections. A series of sections spaced 200 µM apart and spanning the striatum was stained for neuronal nuclei using metal enhanced DAB detection and mounted onto slides for imaging (**Figure 1A**).

**Figure 1.**
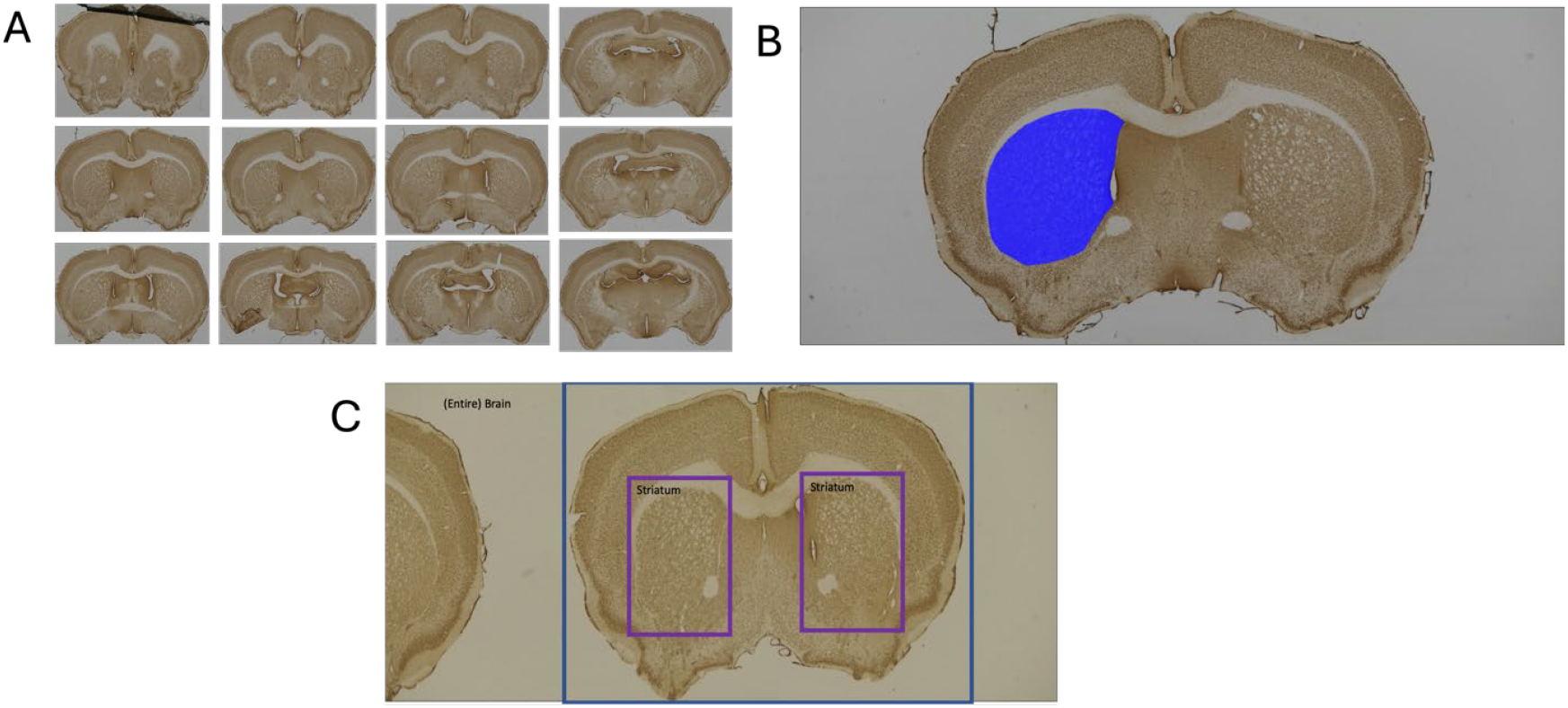
Imaged coronal cross-sections were used for training and testing. (**A**) An evenly spaced series of 12-month-old Hu97/18 humanized HD model mouse imaged coronal brain cross-sections spanning the striatum and stained for neuronal nuclei. (**B**) Example of one hemisphere of the wanted brain region, in this case the striata, being segmented. (**C**) Boxed regions of interest around the entire cross-section and both striata.

Both the left and right striatum from each section were traced. Based on those tracings, a determination of the cross-sectional area can be made. As such, it is those tracings that we want to be automated and more accurately completed through our program.

The way in which manual tracing can be automated is through image segmentation. Image segmentation refers to explicitly outlining each distinct feature of an image. As an example, given an image depicting the cross-section of an HD mouse brain, a successful segmentation would explicitly outline the striata, and any other desired region(s) (**Figure 1B**). There have been multiple advancements made in the way of image segmentation^6^. We have leveraged the strengths of multiple AI algorithms and an algorithm based on TDA to compile a novel Neuropathology Assessment Tool (NAT).

### Creation of the AI Model

The NAT consists of three major parts. The first of which is the Faster Region Based Convolutional Neural Network^7^ (R-CNN). The idea behind the Faster R-CNN is to “crop” out the wanted regions of an image before segmentation, as shown above (**Figure 1C**). This has been implemented for two reasons. The first of which is accuracy. Within our preliminary work, we found that segmentation algorithms tend to work better when they are primarily shown what they are meant to segment. A good way to think about this is that if our segmentation algorithms were given an image of the entire brain, while it is not likely to happen, there is still a chance that it could classify a pixel inside the cortex as being part of the striatum. If our AI model is only able to look at a cropped view of the striatum, this potential inaccuracy is eliminated. The second is that the goal of this technology is distribution. The smaller the image that the AI must process, the easier it is on a computer. The Faster R-CNN can handle larger images based on the way it was designed, it is not nearly as computationally intensive as the rest of the NAT. Therefore, cropping the image down allows not only a more accurate but faster evaluation of the brain region wanting to be segmented.

One of our main goals was to maximize the accuracy of image segmentation. Some of the most recent advancements made in the way of accuracy have been from Transformers^8^ which capitalize on the self-attention techniques. Self-attention is designed to look at all the information given, and selectively attend to the important information. Originally, we planned to use an adaptation we created of the Visual Transformer (ViT)^9^ to create a classification grid of the image, separating it based on striatum or not (**Figure 2A**). The way this adaptation worked was by gridding the image into patches and then applying self-attention over each patch, which we deemed as patched attention (**Figure 2B**).

**Figure 2.**
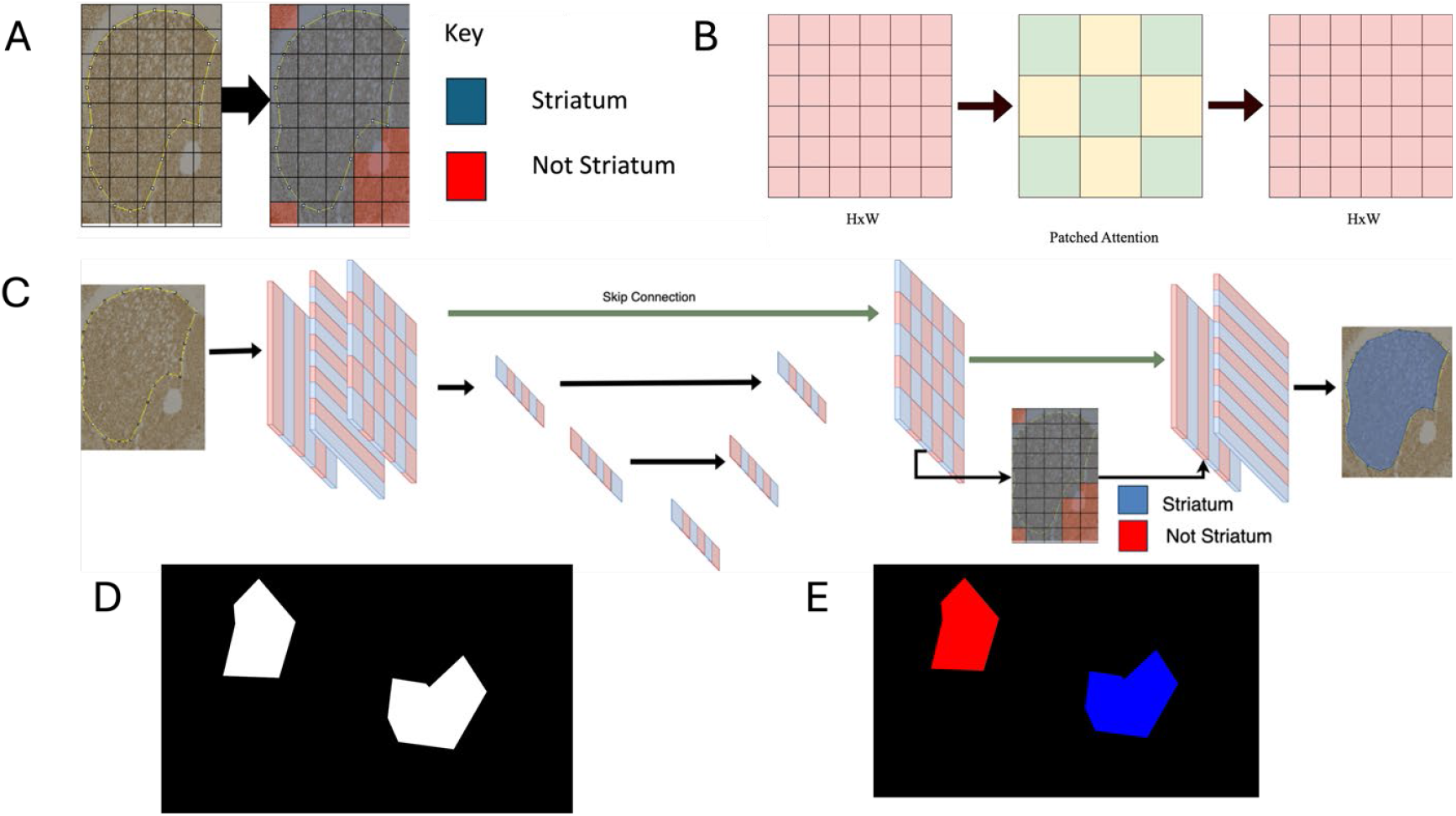
The creation of our tool spans the use of artificial intelligence and topological data analysis. (**A**) Our gridded classification model determines which patches of an image contain the wanted brain region of interest or not. (**B**) Our patched attention mechanism breaks an image into patches and performs self-attention on each patch. (**C**) The mechanistic overview of the artificial intelligence component of our tool, which is deemed the U* Transformer. (**D**) Distinct shapes are represented in white inside an image. (**E**) The use of discrete path connectedness to find individual shapes is a major aspect of the topological data analysis feature of our tool.

Unfortunately, the ViT adaptation fell short in terms of accuracy. Moreover, there were computational challenges when integrating our adaptation into the NAT framework.

Nevertheless, we found the concept of gridded classification and its potential for improving accuracy to warrant further investigation.

During this further investigation, we developed that was deemed the U* Transformer, which forms the AI model of our NAT. This tool capitalizes on the transformers^8^, axial attention^10^, as well as our own approaches including patched attention (**Figure 2C**).

### Training our U* Transformer

Upon the successful completion of the U* Transformer we transitioned into the training of the AI, and validation of the segmented images. We used the dataset created from the Hu97/18 humanized transgenic mouse model to train the U* Transformer. We later used the dataset created from the Q175FDN HD model mice to assess the accuracy of our model. The way the training process happens through a continuous refinement of each function within our models’ parameters. This iterative process is the foundation for the ability of AI to “learn”. Once training is completed, the validation brain set, distinct from the set used to train the AI, was used to compare the accuracy of our algorithm against brains it has never processed before.

While the design of our U* Transformer is a vitally important aspect, execution and training process are equally crucial. Originally, despite the computational advances made, the training process of our U* Transformer, which is what enables our AI to learn to recognize and trace the striatum, was still too computationally expensive for an individual computer.

Consequently, we employed the supercomputer at the UCF Newton Advanced Research Computing Center. Access to this computer allowed the automated training of our AI to be completed within only a few days. This accelerated training period provided valuable insights, allowing the advancement of the U* Transformer based on what was observed.

Another crucial aspect of training our U* Transformer was pre-processing our images. Looking at the actual shape of the striatum, it can be noticed that there are 3 fairly distinct shapes. The first mimicking a rib-eye shape, transitioning into an ovular shape, and finally a more curved kidney bean shape. For this reason, we decided to cluster the images into three distinct subsets, and train three distinct U* Transformer models based on this clustering. We decided to use the Structural Similarity Index Matrix (SSIM)^11^ to perform this clustering for us, instead of potentially relying on a third AI model. The next thing that we did was perform histogram equalization on the images^12^. Histogram equalization is particularly useful for images that have been stained and imaged under microscopy, as it enhances contrast by redistributing the intensity values, making structures within the tissue more distinguishable. This step ensures more consistent feature extraction across varying imaging conditions. Additionally, the validation set was clustered by comparing its SSIM to that of the training dataset, allowing us to match the validation images more closely with their respective clusters. This clustered validation set was then used to validate the performance of the U* Transformer models, after they were trained on the original clustered dataset.

### The refinement of our NAT

The culmination of our NAT lies in the refinement algorithm we used that employs many of the concepts behind topological data analysis (TDA). TDA offers a powerful method to analyze complex high-dimensional data, using concepts from topology^13^. Topology, which can be employed to study the properties of geometric objects that remain unchanged through continuous deformations such as stretching or bending^14^, provides a way to understand the intrinsic shape of data beyond traditional methods, such as AI. TDA allows us to capture the relationships between data points, identifying patterns and structures that are often missed by other methods, making it especially useful for problems where the geometry and connectivity of the data play a crucial role.

Specifically focusing on how TDA can be used with images, one of the fundamental aspects is fitting the data within our image into a metric space and building simplicial complexes from our data.

When defining a metric space (*M, d*) upon a segmented image in 2D space with an image of size *H* × *W H, W* ⊂ ℕ where *M* ⊆ *H* × *W*, such that ∀ *p*ϵ*M, p* = (*x, y*), *x* ϵ *H, y* ϵ *W*, the distance function can be defined as ∀ *p*_1_, *p*_2_ϵ*M*, such that 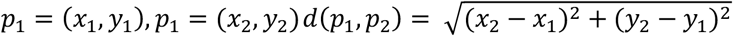.Using this, a form of discrete path connectedness can be used to define two connected spaces. Specifically, in the context of images, subsets can be defined as connected based on either 4-connectivity or 8-connectivity. This means that a subset *I* ⊆ M is connected if ∀*p*1, *p*2 ∈ *I*, ∃{*p*1, *p*3, …, *pn, p*2} ⊆ *I* Such that 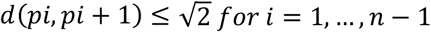.**Figure 2D** shows all elements of *M* such that each pixel in white represents an element in *M* based on its (*x, y*) positioning in the image. When using discrete path connectedness, we find there are two discrete path connected subsets of *M* represented by different colors (**Figure 2E**).

The distance formula is a fundamental tool in defining simplices, which are basic building blocks in TDA. Using the same distance formula for our metric space (*M, d*) defined above, a simplex can be defined by forming connections between points that are within a specified radius *r*. For example, a 0-simplex corresponds to a single point, a 1-simplex is formed by connecting two points that are within the distance *r*, a 2-simplex is created by connecting three points to form a triangle if all pairwise distances are less than *r*, and higher-dimensional simplices are formed similarly. This approach allows for the construction of simplicial complexes, which are collections of simplices that capture the spatial relationships between data points in a way that reflects their underlying topological structure.

Using these concepts from TDA, we created a final refinement algorithm for our NAT. Again, the anatomy of the striatum is heavily considered. The striatum is a singular piece, for this reason, we start by finding all discrete connected spaces, using the same method as described above, followed by only processing the largest one. This ensures that our final output is as close to anatomically correct as possible. From there, each pixel that is classified as striatum is treated as its own distinct data point, and simplicial complexes are formed around each one. The radius for the simplicial complexes stops increasing once all points are connected by a simplex. From there, each actual simplex is filled in. Anything that is within the area outlined by the U* Transformer, and is within the main discrete connected area, is filled in.

## Results

Upon completion of our NAT design and training, there were multiple endpoints which needed to be measured in order to determine success. The first of which being the ability of the RCNN to accurately extract the striatum from each image. We found that the Faster-RCNN was able to crop out a region that entirely contained the wanted striata (**Figure 3A**). This is quantitatively recapitulated by the extracted bounds within each image (**Figure 3B**).

**Figure 3.**
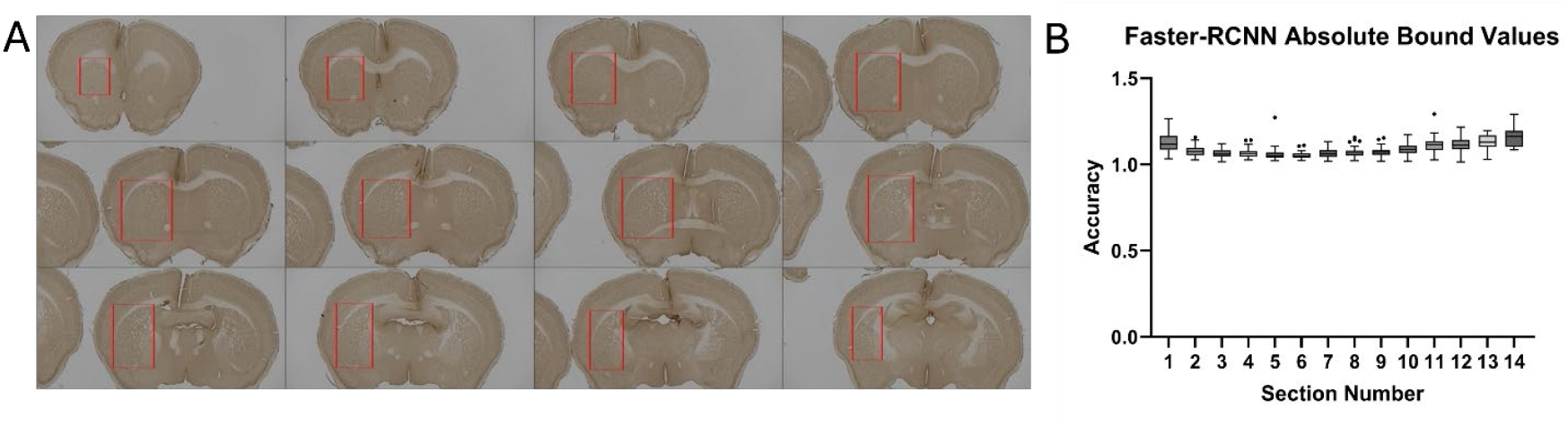
Using the Faster-RCNN the wanted brain region can be entirely cropped out of the imaged cross-section. (**A**) Our use of the Faster-RCNN is able to put bounding boxes in red around the left striata from each imaged cross-section. (**B**) A box and whisker plot representing the percentage of striatum obtained for each anterior-posterior section number when using the Faster-RCNN (N= 10-55). In all cases, all the striatum was obtained, and there was also additional surrounding brain within each bounding box.

Once we found the Faster-RCNN was able to extract the striatum from imaged cross sections successfully, we evaluated the accuracy of our segmentation model, the U* Transformer. We evaluated the accuracy of the model on a metric that was pixel by pixel. This metric determined whether each pixel was correctly identified as part of the striatum or not, focusing specifically on the extracted regions rather than the entire cross-section. Depending on anterior-posterior position within the brain, the striatum has three distinct shape clusters (**Figure 4A**), for which we designed three different U* Transformers; one to handle each cluster. The validation set included 1,368 images that the U* Transformers had not been trained on, and were extracted from the Faster-RCNN (**Figure 4B**). Qualitative results, including representative images, visually demonstrate the segmentation performance and accuracy of the model (**Figure 4C**).

**Figure 4.**
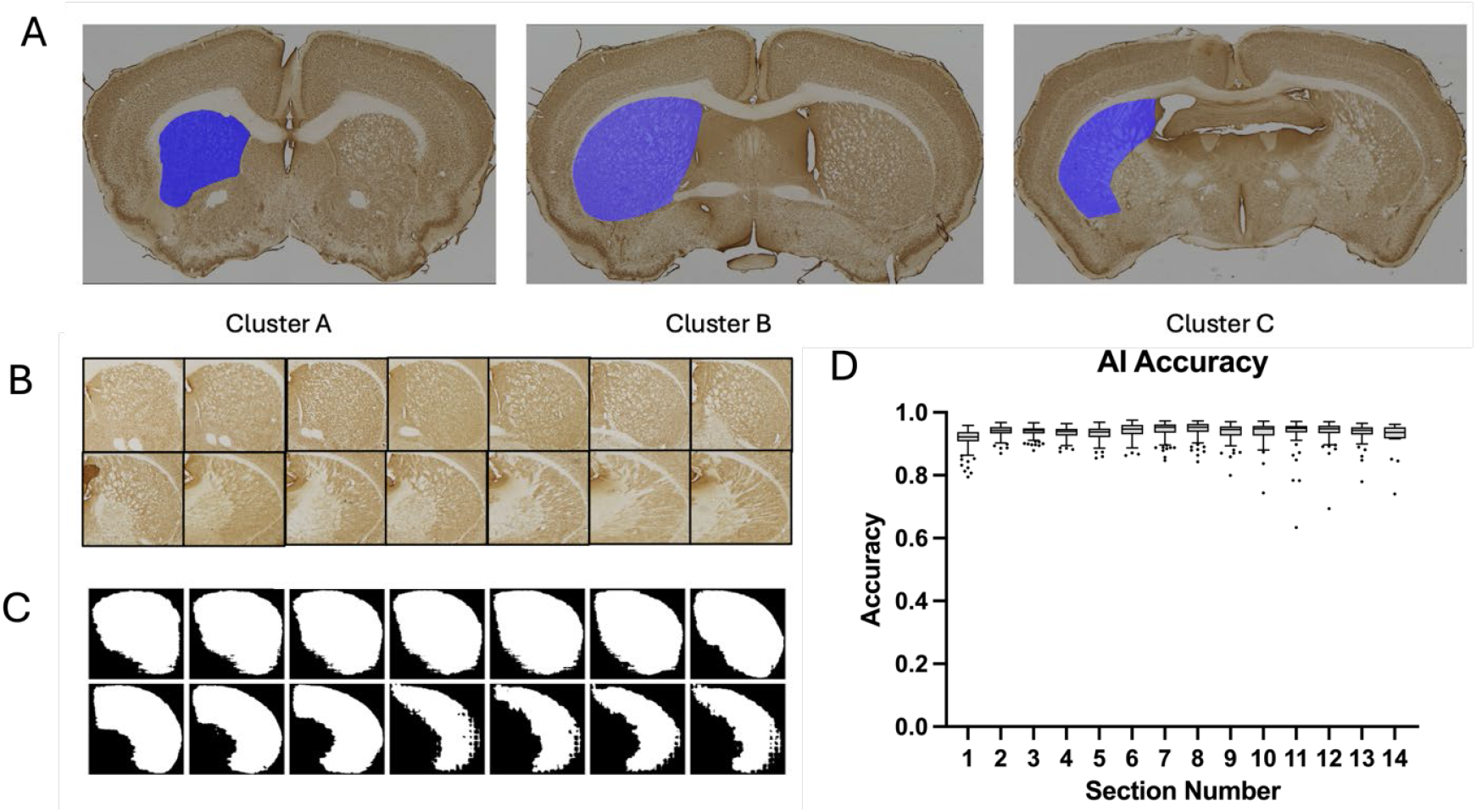
The U* Transformer is able to accurately segment the wanted brain region. (**A**) The shape of the striatum in coronal brain sections has three distinct structural clusters depending on anterior-posterior position within the brain. An individual U* Transformer copy is used for each cluster, increasing the accuracy of our tool. (**B**) An example of the output of the Faster-RCNN, which is then run through the U* Transformer. (**C**) Outputs from the U* Transformer after being given the images obtained from the Faster-RCNN. (**D**) A box and whisker plot representing the accuracy of the U* Transformer based on the cross-section number (N = 20-110), averaging an accuracy of 93.6%.

Quantitatively, the U* Transformer was able to achieve an average accuracy of 93.6% (**Figure 4D**).

While quantitatively, there was a high accuracy associated with the output of the AI portion of our NAT, qualitatively, there were still discrepancies. Specifically, there were regions within the striatum that were not classified as striatum, which do not recapitulate the biological nature of the striatum as a single structure. Additionally, there were isolated patches outside of the main striatum shape that were classified as striatum. For this reason, we developed a novel refinement algorithm based on the concepts behind topological data analysis (TDA)^13^. The TDA refinement algorithm works by finding the main shape in each output, which is the striatum, and filling that shape while eliminating the isolated patches. We applied the refinement algorithm to each segmentation prediction generated by the U* Transformer (**Figure 5A**). We observed a noticeable visual improvement in the segmentation quality after applying the refinement algorithm (**Figure 5B**). Additionally, our algorithm was able to raise the accuracy of the segmented images to an average of 93.8% (**Figure 5C**).

**Figure 5.**
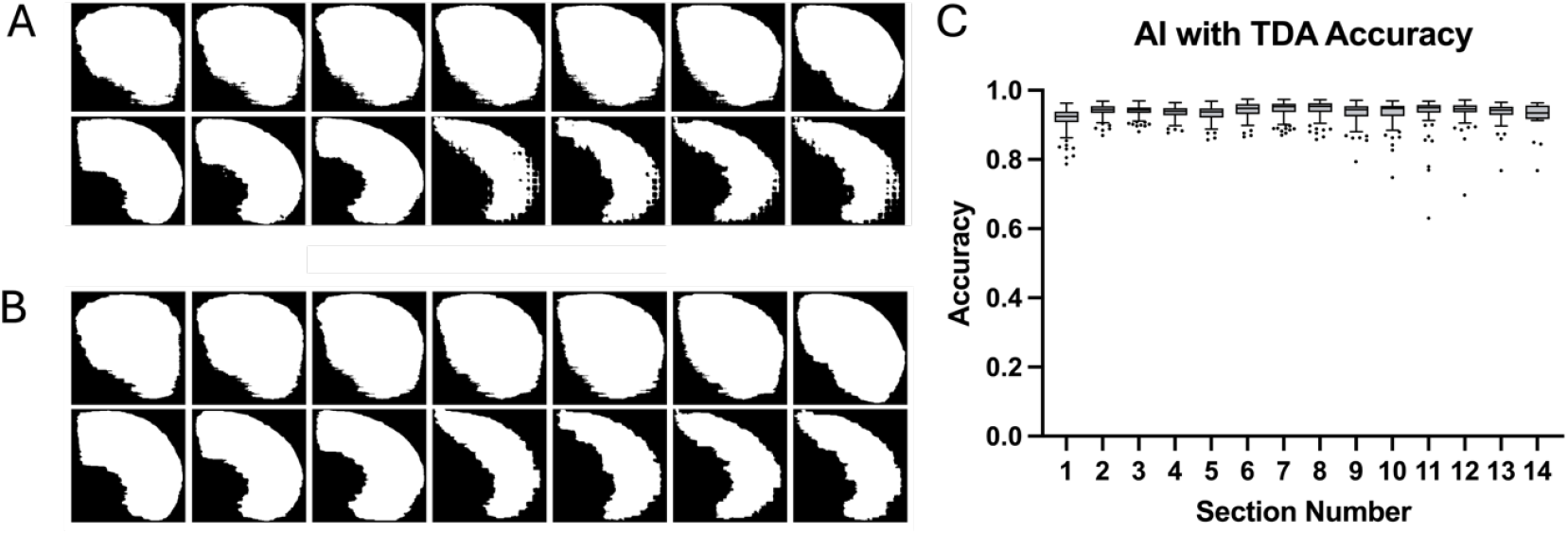
A refinement algorithm based on topological data analysis raises the accuracy of our tool. (**A**) An example of the output of the U* Transformer, which is then run through the refinement algorithm. (**B**) Qualitative results of the refinement algorithm after being given outputs obtained from the U* Transformer. (**C**) A box and whisker plot representing the accuracy of the U* Transformer with the TDA refinement algorithm based on the cross-section number (N=110-20), averaging an accuracy of 93.8%.

After the completion of our NAT design and training, we needed to evaluate its ability to assess neuropathology. Specifically, we used the NAT to measure striatal volumes in Q175FDN heterozygous (HET) HD model mice and wild-type (WT) littermate controls, aiming to determine if it could reliably and accurately detect previously reported genotypic differences^5^, with a higher efficiency compared to manual assessments, while maintaining strong agreement with manual measurements and significantly lower inter-group variability.

Analysis comparing the volumes from NAT and manual analysis showed no significant difference (p=0.28) (**Figure 6A)**. Overall, there was a 2.7% difference when comparing volumes from manual and NAT assessments. As is consistent with historical data, there was a significant difference in striatal volume between Q175FDN HET and WT mice when analyzed using both manual (p = 6E-6) and NAT (p = 9E-6) assessments (**Figure 6B**). The manual method detected 11.24% atrophy, while the NAT detected 10.37% atrophy. There was no significant difference when comparing WT to WT groups (p = 0.28). There was no significant difference when comparing HD to HD groups (p = 0.46). When using a simple linear regression model to analyze the relationship between manual and NAT volumetric data. The model has an R-squared value of 0.945, indicating that 94.5% of the variance in the volumes generated by the NAT is explained by the Manual tracings. The p-value for the model suggests the relationship is significant (P = 4.3E-35). The 95% confidence interval of the slope is [0.98, 1.12], and for the intercept is [-0.63, - 0.43] further confirming the strength of the correlation (**Figure 6C**).

**Figure 6.**
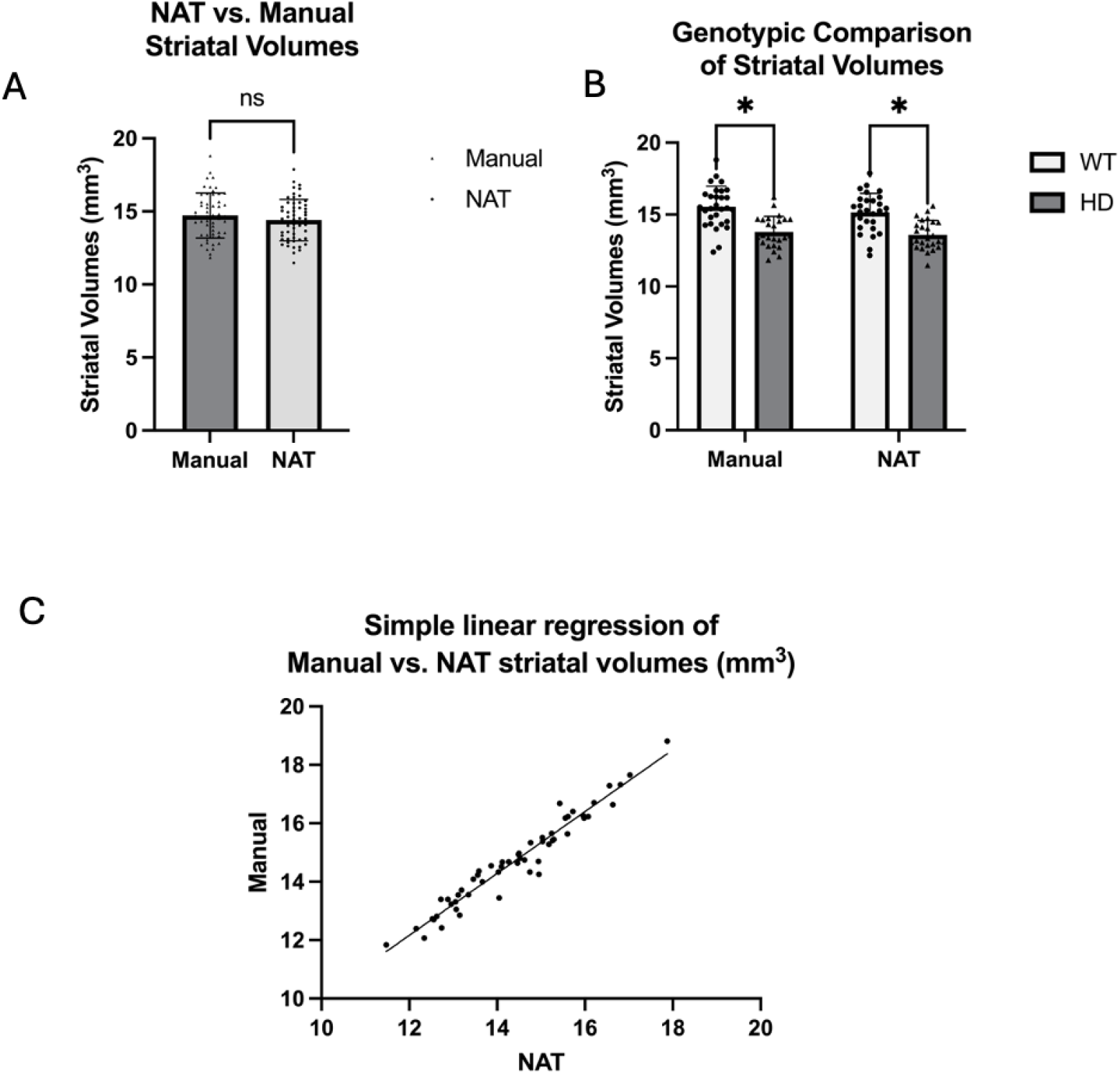
Automated NAT successfully recapitulates manual striatal volume analysis. (**A**) Comparison of striatal volumes assessed manually (n=55) and by the NAT (n=55) using a Student’s t-test showed no significant difference (p=0.28). (**B**) Pairwise t-tests displayed a significant difference in striatal volumes between HD (n=27) and WT (n=28) groups, in both manual (p=6E-6) and NAT (p=9E-6) volumetric analysis. No significant differences were found when comparing HD to HD (p=0.46) or WT to WT (p=0.28) groups. (**C**) A simple linear regression model displaying the relationship between manual (n=55) and NAT (n=55) volumetric data had an R-squared value of 0.945, indicating that 94.5% of the variance in NAT-generated volumes is explained by manual tracings. The model is significant (p = 4.3E-35), with a 95% confidence interval for the slope of [0.98, 1.12] and for the intercept of [-0.63, -0.43].

## Discussion

The integration of the Faster R-CNN, U* Transformer, and our TDA-based refinement algorithm has led to the development of our Neuropathology Assessment Tool (NAT), a tool specifically designed for evaluating neuropathology in preclinical models. Our results demonstrate that the NAT can accurately assess striatal volumes and detect subtle neuropathological changes in Q175FDN HD model mice, even when applied to datasets, and other models of mice, that it was not originally trained on. This highlights the generalizability of our tool across different genotypes, which is crucial for its application.

One of the major strengths of our NAT lies in its efficiency compared to traditional manual methods. Our analyses revealed a high degree of agreement between the NAT and manual measurements, with no significant differences between the two methods and an overall difference of 2.7% in volume assessments. Importantly, the NAT was able to maintain sensitivity in detecting genotype-specific differences in striatal atrophy, showing a significant difference in striatal volume between Q175FDN HET and WT mice, consistent with historical data^5^. These findings suggest that the NAT can serve as a reliable alternative to labor-intensive manual stereological methods while reducing the time and resources required for volumetric analysis.

Moreover, the linear regression analysis between manual and NAT measurements demonstrated a strong correlation with an R-squared value of 0.945, indicating that over 94% of the variability in manual tracing-derived volumes can be explained by the NAT. The consistency in this relationship, with a slope close to unity and a tight confidence interval, reinforces the robustness of the NAT in producing accurate volumetric assessments. This high correlation suggests that the NAT poses a potential to replacing manual methods without compromising the precision and reliability of measurements.

We are actively working to further demonstrate the generalizability of our NAT across different mouse strains and staining methods. This approach aims to validate the tool’s versatility, ensuring that our NAT can maintain high accuracy and reliability across various staining techniques and genetic backgrounds, ultimately supporting its applicability. In addition, applying the NAT to other brain regions will require iterative tracing and additional training to accurately capture features specific to other regions. Adding this additional support for other brain regions is imperative for use in a wide range of neurodegenerative diseases, and to demonstrate the generalizability of the NAT. The ability to assess neuropathological endpoints reliably, such as striatal atrophy, is critical in preclinical trials. Interventions that delay or halt atrophy are most likely to be disease-modifying, highlighting the impact of the NAT for accelerating the identification and testing of such potential treatments.

The culmination of the NAT’s success offers the potential for an increased efficiency of preclinical evaluation of neuropathology, enabling a more rapid testing of experimental therapies. This increased throughput holds a potential to accelerate drug discovery efforts for HD and other intractable neurodegenerative diseases, providing a more streamlined and effective path to identifying promising treatments.

## Acknowledgments

The authors would like to acknowledge the researchers at the UCF Newton Advanced Research Computing Center without their help, our project would never have advanced the way it did. We would also like to thank the HDSA for the ability to work on this project through the HDSA Donald King Fellowship.

## Funding

Support for A.L.S. provided by The National Institutes of Neurological Disorders and Stroke (R01NS116099). Support for S.M. provided by The Huntington’s Disease Society of America Donald A. King Fellowship and The University of Central Florida Office of Undergraduate Research Grant.

## Notes

### Competing Interest Statement

The authors have declared no competing interest.

